## Supplementary Materials for "The neural correlates of shared and individual experience"

### Supplementary Material 1

***Temporal autocorrelation bias in correlations between neural and behavioural time-by-time similarity matrix.***

We aimed to detect correspondence between the subjective (i.e., suspense) and neural dynamics via the time-by-time similarity matrix (TTM see figure 1C, 1E and 1G; and methods).

Given that the subjective and neural TTMs were acquired from different cohorts, and that we primarily wanted to investigate any correspondence between the two, we obtained a TTM for each sample (subjective and neural), by getting the average and the coefficient of variance across subjects. To estimate a significance value of such a correspondence we used a permutation analysis by which we shuffled the columns of the subjective TTM, corresponding to the timepoints of the story 10000 times (e.g., Kriegeskorte, 2015). This yielded a null distribution of correlation values arising out of TTM that have had their timepoints shuffled.

However, TTM have a characteristic degree of autocorrelation (see values along to diagonal fig. 1C and E). By autocorrelation, we mean the increased similarity between closely spaced time points, which are shown along the diagonal of the TTMs. It is expected that timepoints that close to each other will tendentially be more similar, both in terms of a stream of mental contents and in terms of neural states. Although this autocorrelational structure is relevant data to the present project (e.g., showing sharp transitions or a “lingering” in a state), this structure will increase correlation values between TTMs. However, of importance here is that a permuted TTM will not have this autocorrelational structure, therefore inflating p-values calculated via such a permutation analysis. Therefore, to avoid a bias in significance probabilities we removed autocorrelational values for the permutation analyses.

To establish how many autocorrelational values to remove for the permutation analysis we plotted the similarity between the subjective and neural TTM as a function of the amount of removed autocorrelation (proximal timepoint similarities) values (fig. S1A for average TTM and S1B for coefficient of variance TTMs).  This was done by averaging across all networks (n=18) for all participants in the awake condition. By getting the first derivative of the curves we have an illustration of how similarity decreased as a function of removed autocorrelation (fig. 1A and 1B), we found two points in which the similarity “stabilised”, indicating decreased influence of autocorrelation (Fig. S1C for average and Fig S1D for coefficient of variance). The similarities stabilised by removing around 10 and 24 proximal time similarity values. These removed values were adopted for the permutation analyses. In the main text we present the correlation (Tau-A; $\tau_A$) values between the full (no autocorrelation values removed) but the p-values of the permutation analyses of the TTM with 10 proximal time similarity values removed. Full results can be found in S2 (average TTM correspondence) and S3 (coefficient of variance TTM correspondence).

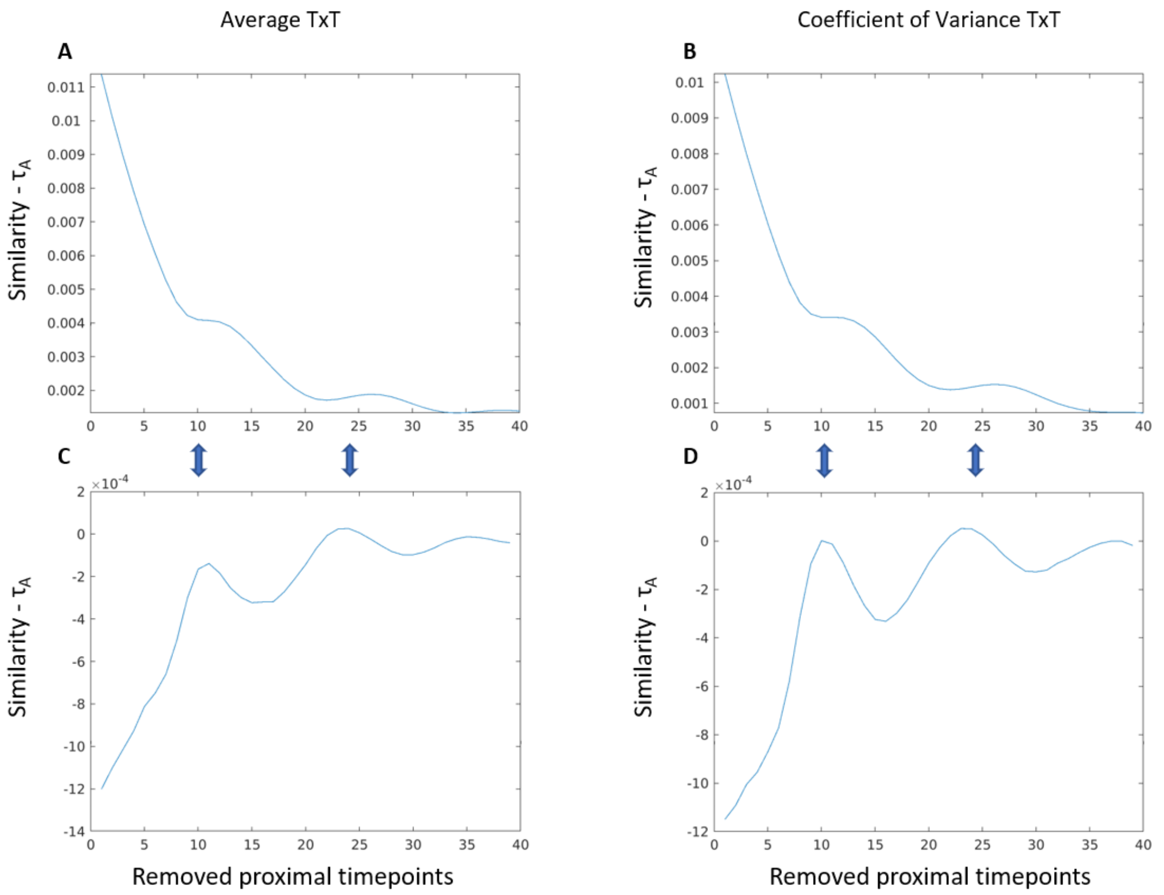


**Figure S1. Similarity (Kendal's $\tau_A$) as a function of removed autocorrelation.**

A and B show the diminishing similarity ($\tau_A$) as a function of removed proximal timepoint similarities (i.e., autocorrelation, values along the diagonal in TTM, e.g., see fig. 1C) for the similarity between subjective and neural average TTMs and Coefficient of Variance TTMs respectively. Figures C and D show the first derivative of A and B respectively revealing two points at which similarities stabilised despite increasing number of autocorrelation values removed. Blue bi-directional arrows indicate the points chosen (i.e., 10 and 24) for the permutation analyses in accordance with the stabilisation of similarities as a function of removed autocorrelation.

### Supplementary Material 2

#### Similarity between neural and subjective average TTM for deconvolved and lagged BOLD data for all networks.

We sought to detect the correspondence between the average neural and subjective dynamics (i.e., TTM) by various means. As shown above we calculated p-values from the permutation analyses (n=10000) by removing 10 and 24 proximal timepoints. We accounted for hemodynamic delay via deconvolving the BOLD timeseries (see methods; (Wu et al., 2021, 2013), the results for which are presented in the main text (Full results presented in table S1). To assess robustness of results we ran a confirmatory analysis where we introduce a delay in the BOLD signal (See Table S2) (Jääskeläinen et al., 2008; Khosla et al., 2021; Nishimoto et al., 2011), corresponding to the peak of the canonical hemodynamic response function (i.e., 6s) as implemented in SPM (https://www.fil.ion.ucl.ac.uk/spm/). Note, uncorrected p-values are presented here (corrected values presented in the main text).

**Table S1. Deconvolved BOLD: Correlation ($\tau_A$) and p-values for correspondence between neural and subjective average TTM.**

Correlation (Tau-A) and permutation test values between the deconvolved neural TTM of each network (600 and 800 granularities) and the subjective suspense TTM averaged across participants. Presented are correlation values between the full subjective and neural TTMs as well as correlation and p values of TTM correspondence having removed 10 and 24 proximal timepoint similarities (see S1). These results are presented visually in figure S2.

| Network Names | $\tau_A$-full | $\tau_A$-10 | P-10 | $\tau_A$-24 | P-24 |
|---|---|---|---|---|---|
| DefaultA 600 | 0.32 | 0.21 | <0.0001 | 0.15 | <0.0001 |
| DefaultA 800 | 0.33 | 0.22 | <0.0001 | 0.16 | <0.0001 |
| DefaultB 600 | 0.22 | 0.10 | 0.0013 | 0.10 | 0.0178 |
| DefaultB 800 | 0.19 | 0.05 | 0.0172 | 0.03 | 0.322 |
| DefaultC 600 | 0.17 | 0.02 | 0.1857 | 0.03 | 0.1457 |
| DefaultC 800 | 0.16 | 0.01 | 0.6003 | 0.01 | 0.5959 |
| TempPar 600 | 0.07 | -0.10 | 0.0225 | -0.09 | 0.0086 |
| TempPar 800 | 0.11 | -0.04 | 0.3655 | -0.06 | 0.1167 |
| ContA 600 | 0.21 | 0.06 | 0.0041 | 0.03 | 0.2149 |
| ContA 800 | 0.23 | 0.09 | <0.0001 | 0.06 | 0.0233 |
| ContB 600 | 0.17 | 0.02 | 0.442 | -0.04 | 0.3239 |
| ContB 800 | 0.23 | 0.09 | <0.0001 | 0.03 | 0.0743 |
| ContC 600 | 0.21 | 0.08 | 0.0357 | 0.03 | 0.1304 |
| ContC 800 | 0.27 | 0.16 | <0.0001 | 0.14 | 0.0002 |
| LimbicA 600 | 0.18 | 0.04 | 0.2642 | 0.04 | 0.2536 |
| LimbicA 800 | 0.16 | 0.01 | 0.6969 | 0.00 | 0.841 |
| LimbicB 600 | 0.13 | -0.04 | 0.566 | -0.12 | 0.0038 |
| LimbicB 800 | 0.18 | 0.02 | 0.2806 | -0.05 | 0.1846 |
| SalVentAttnA 600 | 0.27 | 0.15 | <0.0001 | 0.12 | 0.0008 |
| SalVentAttnA 800 | 0.29 | 0.17 | <0.0001 | 0.15 | 0.0009 |
| SalVentAttnB 600 | 0.16 | 0.00 | 0.8074 | -0.01 | 0.6404 |
| SalVentAttnB 800 | 0.18 | 0.02 | 0.0855 | 0.00 | 0.8317 |
| DorsAttnA 600 | 0.17 | 0.02 | 0.144 | 0.00 | 0.8574 |
| DorsAttnA 800 | 0.17 | 0.03 | 0.055 | -0.02 | 0.4072 |
| DorsAttnB 600 | 0.19 | 0.04 | 0.0554 | -0.04 | 0.1405 |
| DorsAttnB 800 | 0.21 | 0.07 | 0.003 | -0.02 | 0.4702 |
| SomMotA 600 | 0.24 | 0.10 | 0.0004 | 0.04 | 0.1557 |
| SomMotA 800 | 0.26 | 0.13 | 0.0003 | 0.07 | 0.0653 |
| Auditory 600 | 0.21 | 0.07 | <0.0001 | 0.01 | 0.5789 |
| Auditory 800 | 0.22 | 0.07 | <0.0001 | 0.01 | 0.452 |
| VisPeri 600 | 0.21 | 0.07 | 0.0445 | 0.02 | 0.3369 |
| VisPeri 800 | 0.23 | 0.10 | 0.0056 | 0.06 | 0.0638 |
| VisCent 600 | 0.21 | 0.09 | 0.0459 | 0.06 | 0.1183 |
| VisCent 800 | 0.21 | 0.08 | 0.045 | 0.06 | 0.1138 |
| SUB 54 | 0.18 | 0.06 | 0.0033 | 0.03 | 0.27 |

**Table S2. Lagged BOLD: Correlation ($\tau_A$) and p-values for correspondence between neural and subjective average TTM.**

Correlation (Tau-A) and permutation test values between the lagged (6s) neural TTM of each network (600 and 800 granularities) and the subjective suspense TTM averaged across participants. Presented are correlation values ($\tau_A$) between the full subjective and neural TTMs as well as correlation and p-values of TTM correspondence having removed 10 and 24 proximal timepoint similarities (see S1).

| Network Names | $\tau_{A}$-full | $\tau_{A}$-10 | P-10 | $\tau_{A}$-24 | P-24 |
|---|---|---|---|---|---|
| DefaultA 600 | 0.34 | 0.23 | <0.0001 | 0.19 | <0.0001 |
| DefaultA 800 | 0.37 | 0.27 | <0.0001 | 0.24 | <0.0001 |
| DefaultB 600 | 0.25 | 0.12 | 0.0004 | 0.12 | 0.0084 |
| DefaultB 800 | 0.24 | 0.11 | <0.0001 | 0.10 | 0.0042 |
| DefaultC 600 | 0.15 | -0.01 | 0.65 | -0.04 | 0.2457 |
| DefaultC 800 | 0.12 | -0.05 | 0.0551 | -0.07 | 0.011 |
| TempPar 600 | 0.03 | -0.15 | 0.005 | -0.17 | 0.0001 |
| TempPar 800 | 0.09 | -0.07 | 0.4823 | -0.10 | 0.0649 |
| ContA 600 | 0.27 | 0.13 | <0.0001 | 0.12 | 0.0025 |
| ContA 800 | 0.27 | 0.14 | <0.0001 | 0.13 | 0.0032 |
| ContB 600 | 0.18 | 0.02 | 0.3211 | -0.04 | 0.1427 |
| ContB 800 | 0.22 | 0.08 | <0.0001 | 0.02 | 0.3009 |
| ContC 600 | 0.26 | 0.13 | 0.0095 | 0.10 | 0.0421 |
| ContC 800 | 0.33 | 0.23 | <0.0001 | 0.22 | <0.0001 |
| LimbicA 600 | 0.12 | -0.05 | 0.0126 | -0.06 | 0.0048 |
| LimbicA 800 | 0.16 | 0.01 | 0.5991 | -0.01 | 0.7094 |
| LimbicB 600 | 0.10 | -0.08 | 0.2697 | -0.16 | <0.0001 |
| LimbicB 800 | 0.13 | -0.05 | 0.7139 | -0.12 | 0.0461 |
| SalVentAttnA 600 | 0.22 | 0.08 | 0.0017 | 0.04 | 0.4022 |
| SalVentAttnA 800 | 0.28 | 0.16 | <0.0001 | 0.11 | 0.0824 |
| SalVentAttnB 600 | 0.21 | 0.06 | 0.0699 | 0.06 | 0.2595 |
| SalVentAttnB 800 | 0.23 | 0.08 | 0.0005 | 0.06 | 0.1141 |
| DorsAttnA 600 | 0.28 | 0.15 | <0.0001 | 0.16 | <0.0001 |
| DorsAttnA 800 | 0.25 | 0.11 | 0.0001 | 0.08 | 0.0177 |
| DorsAttnB 600 | 0.22 | 0.08 | 0.0099 | 0.00 | 0.829 |
| DorsAttnB 800 | 0.26 | 0.13 | 0.0005 | 0.08 | 0.0441 |
| SomMotA 600 | 0.27 | 0.14 | 0.0001 | 0.08 | 0.0245 |
| SomMotA 800 | 0.25 | 0.11 | <0.0001 | 0.04 | 0.128 |
| Auditory 600 | 0.20 | 0.06 | 0.0008 | 0.00 | 0.8884 |
| Auditory 800 | 0.18 | 0.02 | 0.3616 | -0.04 | 0.3477 |
| VisPeri 600 | 0.25 | 0.11 | 0.0022 | 0.07 | 0.0568 |
| VisPeri 800 | 0.23 | 0.09 | 0.0092 | 0.04 | 0.1336 |
| VisCent 600 | 0.20 | 0.06 | 0.0967 | 0.01 | 0.5164 |
| VisCent 800 | 0.25 | 0.12 | 0.0179 | 0.09 | 0.0437 |
| SUB 54 | 0.16 | 0.02 | 0.1687 | 0.00 | 0.99 |

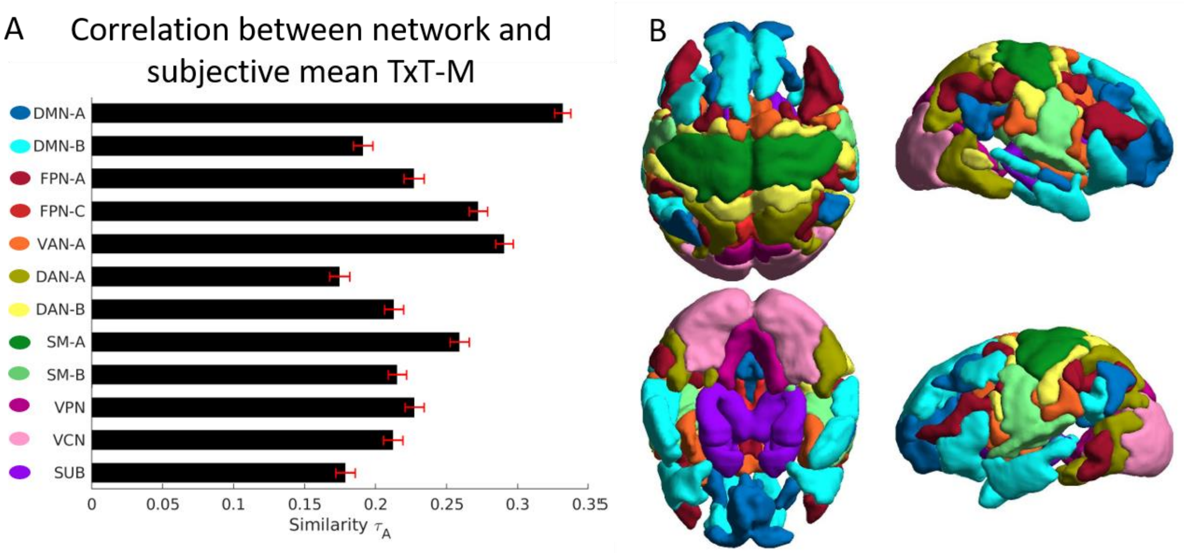


**Figure S2. Correlation between network and subjective mean TTM and representation of networks. – Deconvolved.**

The bar graph (A) shows similarities ($\tau_A$) between the mean subjective and neural TTM for significant networks (see methods and S1). Error bars represent bootstrapped (n=1000) standard errors. (B) The significant networks are spatially represented in B. Colour legend for B can be found next to labels in A. DMN=Default Mode Network; FPN=Frontal Parietal Network; DAN=Dorsal Attention Network; VAN=Ventral Attention Network; SM=SomatoMotor Network (Auditory network in the main text); VCN=Visual Central Network; VPN= Visual Peripheral Network; SUB= Subcortex.

Of note is that although there is large correspondence between the deconvolved and lagged approaches, the convergence is not perfect. Namely the DAN-A and the Subcortex are only significant in the Lagged and Deconvolved data respectively. However it is known that subcortical regions have abnormal haemodynamics compared to the cortex (Lewis et al., 2018; Miletić et al., 2020), potentially explaining the lack of consistency across different analyses. Hence the use of the blind deconvolution toolbox may have been beneficial for the analysis of the subcortex.

As for the DAN-A, it is beyond the scope of this study to investigate the differences between the lag (Jääskeläinen et al., 2008) and the deconvolution (Wu et al., 2013) method, and whether there is an interaction with the instantaneous phase synchrony method used in this study. Suffice it to say that in the deconvolution analysis the correlation values drop drastically (from 0.17 to 0.03) after autocorrelation is maintained whilst in the lagged analysis, they remain reasonably high after autocorrelation is removed (0.11, 0.15). This may suggest that in the lag analysis there is relevant information in the distal parts of the TTMs (i.e., which is not contained in the autocorrelation S1), whilst after deconvolution this is not the case.

Also interesting is that in the TempPar network, which showed a strikingly different behaviour between rest and story conditions (discussed in S4), showed negative correlations between the subjective and neural data once the autocorrelation was remove (Table S1 and S2).

### Supplementary Material 3

**Similarity between neural and subjective coefficient of variance TTM for deconvolved and lagged BOLD data for all networks.**

We sought to investigate whether neural variability between subjects corresponded to subjective variability in temporal domain (as measured by TTM). We calculated the coefficient of variance across participants for the subjective and neural TTM and corresponded them using both the deconvolution and the lagged approach (See summplemntaty material 2 and methods). The coefficient of variance is preferable to standard deviation given the substantial difference in the nature of the subjective data (Likert scale from 1 to 9) compared to the neural data (Pearson correlation values from 1 to -1, between high dimensional connectivity matrices). Similarly, to the correspondence between average TTMs, p-values were calculated from a permutation analysis (n=10000) on TTMs that had the similarities between 10 and 24 proximal timepoints removed (see S1). Note, uncorrected p-values are presented here.

**Table S3. Deconvolved BOLD: Correlation ($\tau_A$) and p-values for correspondence between neural and subjective coefficient of variance TTM.**

Correlation (Tau-A) and permutation test values between the coefficient of variance of the deconvolved neural TTM of each network (600 and 800 granularities) and the coefficient of variance of the subjective suspense TTM. Coefficient of variance was calculated across participants. Presented are correlation values ($\tau_A$) between the full subjective and neural TTMs as well as correlation and p values of TTM correspondence having removed 10 and 24 proximal timepoint similarities (see S1). Significant results are also presented visually in figure S3.

| Network Names | $\tau_{A}$-full | $\tau_{A}$-10 | P-10 | $\tau_{A}$-24 | P-24 |
|---|---|---|---|---|---|
| DefaultA 600 | 0.33 | 0.23 | <0.0001 | 0.2 | <0.0001 |
| DefaultA 800 | 0.35 | 0.27 | <0.0001 | 0.24 | <0.0001 |
| DefaultB 600 | 0.21 | 0.1 | 0.0003 | 0.1 | 0.0099 |
| DefaultB 800 | 0.16 | 0.03 | 0.1174 | 0.01 | 0.7607 |
| DefaultC 600 | 0.14 | 0.01 | 0.7828 | -0.01 | 0.5305 |
| DefaultC 800 | 0.12 | -0.02 | 0.6605 | -0.03 | 0.4418 |
| TempPar 600 | 0.08 | -0.07 | 0.4163 | -0.07 | 0.2521 |
| TempPar 800 | 0.1 | -0.04 | 0.7299 | -0.05 | 0.4296 |
| ContA 600 | 0.19 | 0.06 | 0.0059 | 0.04 | 0.1853 |
| ContA 800 | 0.16 | 0.01 | 0.8964 | -0.02 | 0.4306 |
| ContB 600 | 0.17 | 0.04 | 0.0265 | -0.02 | 0.3431 |
| ContB 800 | 0.2 | 0.08 | 0.0011 | 0.02 | 0.3684 |
| ContC 600 | 0.14 | 0.02 | 0.2818 | -0.01 | 0.7026 |
| ContC 800 | 0.19 | 0.07 | 0.01 | 0.08 | 0.008 |
| LimbicA 600 | 0.13 | 0 | 0.9151 | -0.02 | 0.1982 |
| LimbicA 800 | 0.16 | 0.03 | 0.106 | 0.01 | 0.6325 |
| LimbicB 600 | 0.15 | 0.01 | 0.7975 | -0.07 | 0.1066 |
| LimbicB 800 | 0.16 | 0.02 | 0.2368 | -0.04 | 0.1491 |
| SalVentAttnA 600 | 0.24 | 0.11 | <0.0001 | 0.07 | 0.0228 |
| SalVentAttnA 800 | 0.27 | 0.15 | <0.0001 | 0.11 | 0.0038 |
| SalVentAttnB 600 | 0.16 | 0.02 | 0.3981 | -0.01 | 0.7158 |
| SalVentAttnB 800 | 0.15 | 0 | 0.9206 | -0.04 | 0.1908 |

| | | | | | |
|---|---|---|---|---|---|
| DorsAttnA 600 | 0.16 | 0.02 | 0.3361 | 0 | 0.8665 |
| DorsAttnA 800 | 0.16 | 0.02 | 0.2154 | -0.01 | 0.3958 |
| DorsAttnB 600 | 0.19 | 0.07 | 0.0164 | 0.01 | 0.6998 |
| DorsAttnB 800 | 0.18 | 0.04 | 0.0421 | -0.02 | 0.5124 |
| SomMotA 600 | 0.18 | 0.04 | 0.0068 | -0.02 | 0.3696 |
| SomMotA 800 | 0.18 | 0.04 | 0.0417 | 0 | 0.9298 |
| Auditory 600 | 0.21 | 0.09 | <0.0001 | 0.06 | 0.0029 |
| Auditory 800 | 0.24 | 0.12 | <0.0001 | 0.1 | 0.0002 |
| VisPeri 600 | 0.15 | 0.01 | 0.6151 | -0.03 | 0.2053 |
| VisPeri 800 | 0.2 | 0.07 | 0.0209 | 0.05 | 0.101 |
| VisCent 600 | 0.22 | 0.1 | 0.0046 | 0.08 | 0.0383 |
| VisCent 800 | 0.22 | 0.11 | 0.0056 | 0.1 | 0.0159 |
| SUB 54 | 0.16 | 0.05 | 0.021 | 0.03 | 0.2936 |

**Table S4. Lagged BOLD: Correlation ($\tau_A$) and p-values for correspondence between neural and subjective coefficient of variance TTM.**

Correlation (Tau-A) and permutation test values between the coefficient of variance of the lagged (6s) neural TTM of each network (600 and 800 granularities) and the coefficient of variance of the subjective suspense TTM. Coefficient of variance was calculated across participants. Presented are correlation values between the full subjective and neural TTMs as well as correlation and p-values of TTM correspondence having removed 10 and 24 proximal timepoint similarities (see Supplementary Material 1).

| Network Names | $\tau_A$-full | $\tau_A$-10 | P-10 | $\tau_A$-24 | P-24 |
|---|---|---|---|---|---|
| DefaultA 600 | 0.31 | 0.22 | <0.0001 | 0.21 | <0.0001 |
| DefaultA 800 | 0.33 | 0.24 | <0.0001 | 0.24 | <0.0001 |
| DefaultB 600 | 0.2 | 0.07 | 0.1523 | 0.09 | 0.1468 |
| DefaultB 800 | 0.2 | 0.07 | 0.0848 | 0.08 | 0.065 |
| DefaultC 600 | 0.1 | -0.06 | 0.0749 | -0.09 | 0.0153 |
| DefaultC 800 | 0.09 | -0.07 | 0.0172 | -0.09 | 0.007 |
| TempPar 600 | 0 | -0.18 | <0.0001 | -0.19 | <0.0001 |
| TempPar 800 | 0.07 | -0.08 | 0.0761 | -0.08 | 0.0844 |
| ContA 600 | 0.25 | 0.12 | 0.0006 | 0.1 | 0.0395 |
| ContA 800 | 0.22 | 0.08 | 0.0021 | 0.09 | 0.0102 |
| ContB 600 | 0.21 | 0.08 | 0.0099 | 0.03 | 0.6332 |
| ContB 800 | 0.23 | 0.11 | 0.0008 | 0.07 | 0.1631 |
| ContC 600 | 0.18 | 0.05 | 0.2453 | 0.04 | 0.2209 |
| ContC 800 | 0.26 | 0.15 | 0.0006 | 0.17 | 0.0007 |
| LimbicA 600 | 0.11 | -0.04 | 0.0687 | -0.05 | 0.0224 |
| LimbicA 800 | 0.19 | 0.06 | 0.0034 | 0.07 | 0.0041 |
| LimbicB 600 | 0.14 | -0.02 | 0.7682 | -0.08 | 0.1003 |
| LimbicB 800 | 0.14 | -0.01 | 0.9523 | -0.07 | 0.3489 |
| SalVentAttnA 600 | 0.23 | 0.1 | <0.0001 | 0.08 | 0.0057 |
| SalVentAttnA 800 | 0.27 | 0.15 | <0.0001 | 0.13 | 0.0005 |
| SalVentAttnB 600 | 0.18 | 0.04 | 0.1485 | 0.05 | 0.3263 |
| SalVentAttnB 800 | 0.19 | 0.04 | 0.0735 | 0.03 | 0.552 |
| DorsAttnA 600 | 0.21 | 0.08 | 0.0036 | 0.07 | 0.0568 |

| | | | | | |
|---|---|---|---|---|---|
| DorsAttnA 800 | 0.2 | 0.06 | 0.0306 | 0.04 | 0.3852 |
| DorsAttnB 600 | 0.24 | 0.12 | 0.0014 | 0.08 | 0.1129 |
| DorsAttnB 800 | 0.22 | 0.09 | 0.0142 | 0.05 | 0.1875 |
| SomMotA 600 | 0.23 | 0.09 | <0.0001 | 0.04 | 0.0889 |
| SomMotA 800 | 0.19 | 0.05 | 0.0039 | 0.01 | 0.4947 |
| Auditory 600 | 0.19 | 0.05 | 0.0003 | 0.02 | 0.247 |
| Auditory 800 | 0.19 | 0.05 | 0.0002 | 0.01 | 0.3954 |
| VisPeri 600 | 0.19 | 0.07 | 0.1551 | 0.04 | 0.5825 |
| VisPeri 800 | 0.2 | 0.07 | 0.023 | 0.04 | 0.2727 |
| VisCent 600 | 0.17 | 0.04 | 0.1585 | 0 | 0.8383 |
| VisCent 800 | 0.26 | 0.15 | 0.0001 | 0.13 | 0.0027 |
| SUB 54 | 0.14 | 0.02 | 0.2506 | 0.01 | 0.5783 |

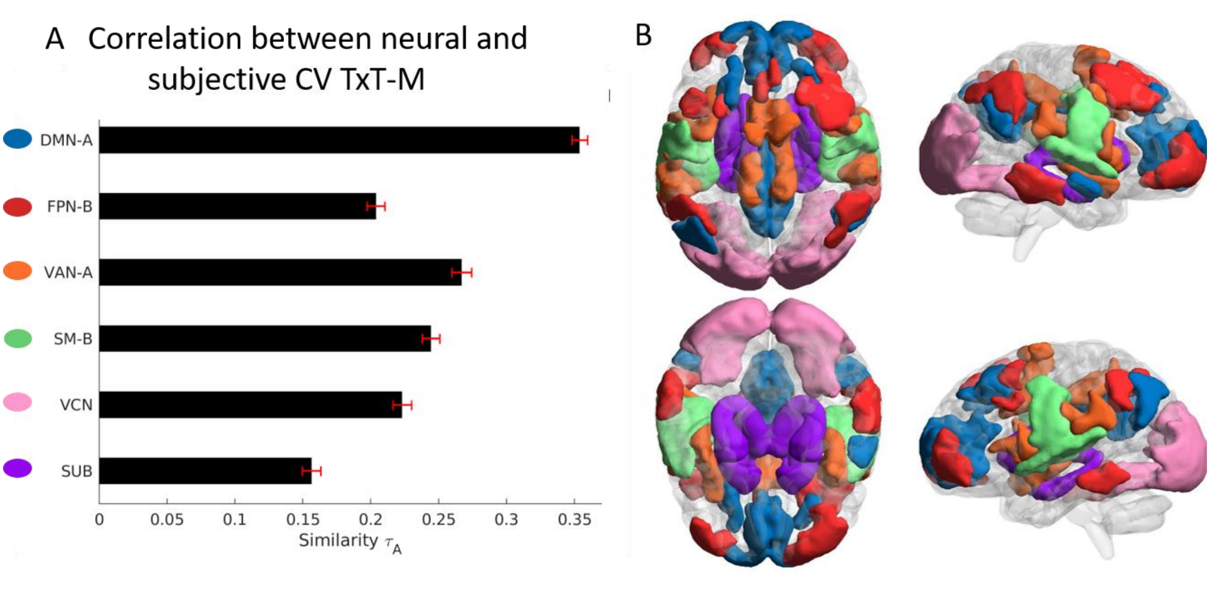


**Figure S3. Correlation between network and subjective coefficient of variance TTM across individuals and representation of networks – Deconvolved.**

The bar graph (A) represents similarity between the coefficient of variance neural and subjective TTM for significant networks. Error bars signify bootstrapped (n=1000) standard errors. The spatial localisations of these networks are presented (B). DMN=Default Mode Network; FPN=Frontal Parietal Network; VAN=Ventral Attention Network; SM=SomatoMotor Network (Auditory network in the main text); VCN=Visual Central Network; VPN= Visual Peripheral Network; SUB= Subcortex.

Once again, there was a good convergence between results. Particularly in the DMN A, SMN A and B, FPN-B, DAN-B.

However, this was not the case for the DMN C, the higher granularity FPN-A, the lower granularity VCN and similarly to the correspondence between average TTMs ( Supplementary material 1) the subcortex and the DAN-A. Interestingly, it seemed that the lagged methods yielded more significant results in the correspondence between the variation across participants in neural and subjective TTMs.

Also, similarly to above the TempPar showed negative correlations between neural and subjective coefficient of variance TTMs once autocorrelation was removed. This, together with the average TTM correspondence analysis (Supplementary material 2) suggests that when the autocorrelation is removed, TempPar varies most when ratings of suspense are stable. This perhaps reflects a higher order, on-going processing that occurs in spite of stable feelings of suspense.

### Supplementary Material 4

**Planned contrasts of inter-subject similarities between awake and deep conditions during story listening for each network.**

We established that there was a significant interaction of inter-subject similarity between the networks (n=18) and the level of anaesthesia (n=3). Therefore, we sought to explore which networks showed higher similarity in consciousness and higher dissimilarity in consciousness (compared to deep anaesthesia), taking this as evidence that these networks would be involved in general and special experiences respectively. We thus ran a Wilcoxon signed rank test on the inter-subject similarities (between each of the 16 participants, thus n=120) of the awake and deep conditions. Here we present the ensuing results for each of the networks across the two granularities.

**Table S6. Wilcoxon signed rank z and p-values of difference between inter-subject similarities in awake and deep anaesthesia conditions.**

| Network Names | Z-value | P-value |
|---|---|---|
| DefaultA 600 | -7.60 | <0.0001 |
| DefaultA 800 | -6.84 | <0.0001 |
| DefaultB 600 | -7.14 | <0.0001 |
| DefaultB 800 | -8.49 | <0.0001 |
| DefaultC 600 | -5.69 | <0.0001 |
| DefaultC 800 | -3.79 | 0.0001 |
| TempPar 600 | -9.08 | <0.0001 |
| TempPar 800 | -8.93 | <0.0001 |
| ContA 600 | 2.84 | 0.0045 |
| ContA 800 | 1.87 | 0.0611 |
| ContB 600 | 1.10 | 0.2714 |
| ContB 800 | -4.36 | <0.0001 |
| ContC 600 | -9.05 | <0.0001 |
| ContC 800 | -9.10 | <0.0001 |
| LimbicA 600 | 7.83 | <0.0001 |
| LimbicA 800 | 8.54 | <0.0001 |
| LimbicB 600 | 1.53 | 0.1249 |
| LimbicB 800 | 3.39 | 0.0007 |
| SalVentAttnA 600 | -3.68 | 0.0002 |
| SalVentAttnA 800 | -3.31 | 0.0009 |
| SalVentAttnB 600 | -4.86 | <0.0001 |
| SalVentAttnB 800 | -5.21 | <0.0001 |
| DorsAttnA 600 | 1.45 | 0.1461 |
| DorsAttnA 800 | 3.14 | 0.0017 |
| DorsAttnB 600 | -0.05 | 0.9624 |
| DorsAttnB 800 | -0.05 | 0.9561 |
| SomMotA 600 | 2.10 | 0.0362 |
| SomMotA 800 | 2.80 | 0.0051 |

| | | |
|---|---|---|
| Auditory 600 | 5.09 | <0.0001 |
| Auditory 800 | 6.62 | <0.0001 |
| VisPeri 600 | 8.27 | <0.0001 |
| VisPeri 800 | 8.16 | <0.0001 |
| VisCent 600 | -7.51 | <0.0001 |
| VisCent 800 | -6.40 | <0.0001 |
| SUB 54 | 8.99 | <0.0001 |

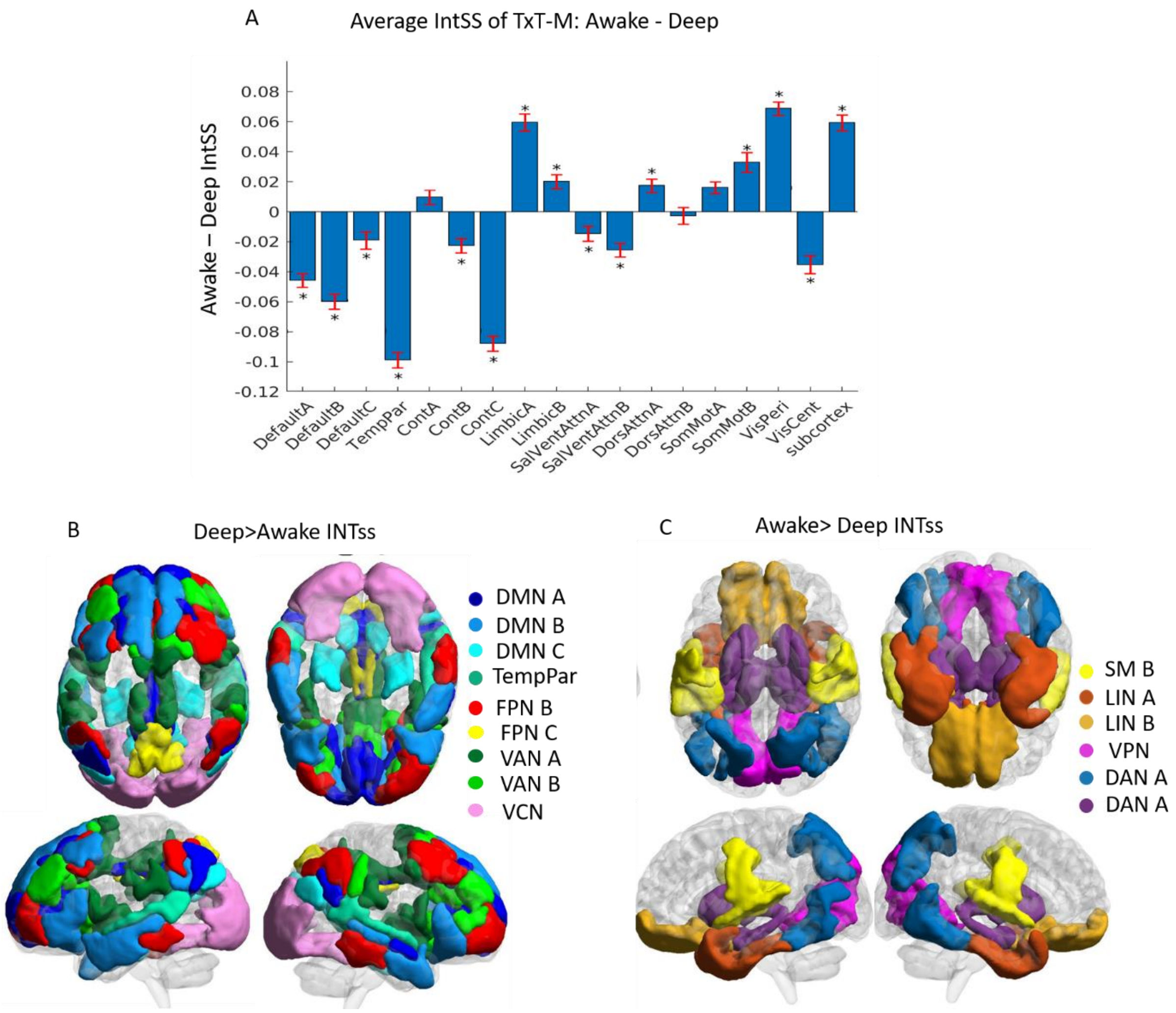


**Figure S5. Inter-subject similarity differences between awake and deep anaesthesia.**

(A) Awake – Deep average inter-subject similarity (IntSS) of TTMs for each network. Asterisks mark FWE corrected significance. Error bars signify bootstrapped (n=1000) standard error of mean difference. Maps showing which systems significantly display more dissimilarity during consciousness (Deep>Awake; B) and more inter-subject similarity (IntSS) during consciousness (Awake>Deep; C). Asterisks indicate FEW corrected significances. Error bars indicate bootstrapped (n=1000) standard error of mean difference.

**Supplementary Material 5**

**3x2 ANOVA for each network. Inter-subject similarities in awake moderate and deep conditions during rest and story listening.**

We wanted to ensure that the proposed "shared and individual" neural correlates of experiences are actually about the contents arising from listening to the story, rather than just a general effect of anaesthesia regardless of story condition. Therefore, we investigated if the inter-subject similarities in the different states of consciousness (awake, moderate anaesthesia, and deep anaesthesia) showed a different pattern of effects between resting state and story listening conditions. We thus built a 3x2 ANOVA (control, moderate and deep anaesthesia, for both rest and story listening) for each network. We particularly focused on the interaction effect in the main text, but here we present results for the two factors (consciousness state and rest vs story task). Of note is that the inter-subject similarities tended to be higher in the resting state condition which is counter intuitive. Upon further investigation, we found that the autocorrelation in the resting condition is typically much higher than the story condition. Perhaps this suggests that the story condition obliges the brain to track the stimuli dynamics more tightly, thus inducing quicker changes over proximal timepoints. This is an interesting finding in its own right, but beyond the scope of the present work.

**Table S5. Values for 3x2 ANOVA for each network. F and P-values presented for interaction between consciousness condition and task vs rest, for task vs rest, and for consciousness condition (awake, moderate, deep anaesthesia).**

| Network Names | F-value Interaction | P-value Interaction | F-value Task vs Rest | P-value Task vs Rest | F-value consciousness condition | F-value consciousness condition |
|---|---|---|---|---|---|---|
| DefaultA 600 | 5.44 | 0.0045 | 6.23 | 0.0128 | 79.19 | <0.0001 |
| DefaultA 800 | 0.15 | 0.8603 | 1.08 | 0.2989 | 116.7 | <0.0001 |
| DefaultB 600 | 31.33 | <0.0001 | 115.39 | <0.0001 | 24.94 | <0.0001 |
| DefaultB 800 | 50.11 | <0.0001 | 91.51 | <0.0001 | 62.75 | <0.0001 |
| DefaultC 600 | 4.96 | 0.0073 | 11.38 | 0.0008 | 31.43 | <0.0001 |
| DefaultC 800 | 12.58 | <0.0001 | 2.2 | 0.1387 | 28.53 | <0.0001 |
| TempPar 600 | 137.33 | <0.0001 | 294.94 | <0.0001 | 22.08 | <0.0001 |
| TempPar 800 | 124.05 | <0.0001 | 365.52 | <0.0001 | 11.07 | <0.0001 |
| ContA 600 | 21.7 | <0.0001 | 19.57 | <0.0001 | 9.43 | 0.0001 |
| ContA 800 | 4.06 | 0.0177 | 13.23 | 0.0003 | 3.5 | 0.0306 |
| ContB 600 | 24.31 | <0.0001 | 27.94 | <0.0001 | 14.81 | <0.0001 |
| ContB 800 | 3.24 | 0.0397 | 6.31 | 0.0122 | 32.44 | <0.0001 |
| ContC 600 | 31.74 | <0.0001 | 82.87 | <0.0001 | 68.85 | <0.0001 |
| ContC 800 | 46.29 | <0.0001 | 93.16 | <0.0001 | 98.87 | <0.0001 |
| LimbicA 600 | 12.42 | <0.0001 | 0.64 | 0.4256 | 136.79 | <0.0001 |
| LimbicA 800 | 35.93 | <0.0001 | 1.25 | 0.2643 | 170.74 | <0.0001 |
| LimbicB 600 | 3.51 | 0.0304 | 60.55 | <0.0001 | 14.58 | <0.0001 |
| LimbicB 800 | 4.98 | 0.0071 | 39.25 | <0.0001 | 31.09 | <0.0001 |
| SalVentAttnA 600 | 1.46 | 0.2322 | 1.11 | 0.2931 | 8.91 | 0.0002 |
| SalVentAttnA 800 | 6.7 | 0.0013 | 5.56 | 0.0186 | 14.6 | <0.0001 |
| SalVentAttnB 600 | 17.85 | <0.0001 | 116.68 | <0.0001 | 3.9 | 0.0206 |
| SalVentAttnB 800 | 9.24 | 0.0001 | 73.92 | <0.0001 | 7.39 | 0.0007 |
| DorsAttnA 600 | 6.77 | 0.0012 | 22.3 | <0.0001 | 14.65 | <0.0001 |
| DorsAttnA 800 | 1.47 | 0.2302 | 30.83 | <0.0001 | 20.43 | <0.0001 |
| DorsAttnB 600 | 2.34 | 0.0973 | 43.28 | <0.0001 | 24.41 | <0.0001 |
| DorsAttnB 800 | 3.61 | 0.0276 | 23.11 | <0.0001 | 35.04 | <0.0001 |

| | | | | | | |
|---|---|---|---|---|---|---|
| SomMotA 600 | 6.16 | 0.0022 | 3.19 | 0.0743 | 30.88 | <0.0001 |
| SomMotA 800 | 13.56 | <0.0001 | 0.04 | 0.8467 | 54.05 | <0.0001 |
| Auditory 600 | 6 | 0.0026 | 6.22 | 0.0128 | 20.47 | <0.0001 |
| Auditory 800 | 7.03 | 0.001 | 1.48 | 0.2237 | 27.63 | <0.0001 |
| VisPeri 600 | 15.3 | <0.0001 | 4.55 | 0.0333 | 53.4 | <0.0001 |
| VisPeri 800 | 1.73 | 0.1781 | 5.35 | 0.021 | 78.33 | <0.0001 |
| VisCent 600 | 15.11 | <0.0001 | 7.91 | 0.0051 | 53.55 | <0.0001 |
| VisCent 800 | 18.45 | <0.0001 | 3.32 | 0.0689 | 28.74 | <0.0001 |
| SUB 54 | 1.72 | 0.1796 | 15.53 | 0.0001 | 109.22 | <0.0001 |

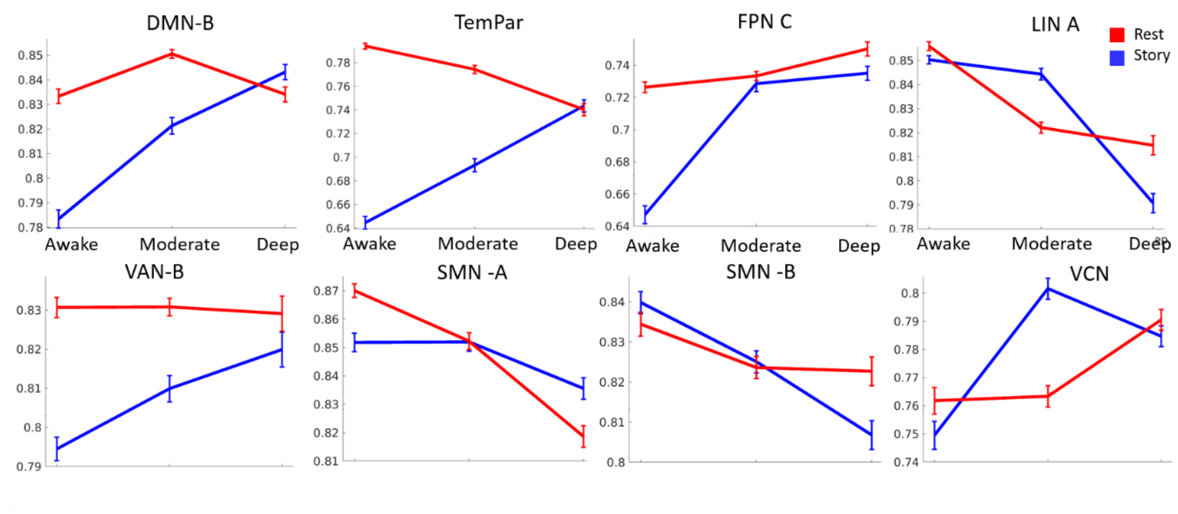


**Figure S4. Rest-story interaction of inter-subject similarities as a function of level of anaesthesia**

Significant interactions of inter-subject similarities from 3x2 ANOVA (Awake Moderate Deep [respectively presented from left to right], for story [Blue] and rest [Red] conditions).

**Supplementary Material 6**

**Inter-subject similarity of TTMs in moderate anaesthesia correlate with reaction times.**

We sought to explore whether individual differences in the effect of moderate anaesthesia (measured via reaction time, See Methods, but also (Deng, Taylor, Owen, Cusack, & Naci, 2020)) would correlate with the amount of inter-subject similarity of neural TTM during the listening of the story. This is motivated by the idea that individual who are most affected by anaesthesia would show a greater breakdown of the "common" (shared) content of experience. We approximated the effect of anaesthesia via a behavioural measure of reaction time, by which individuals would have to respond to a button press to an auditory stimulus. Despite all individuals reported understanding the task, and performing correctly during the awake condition, there were three subjects who timed out, reaching the maximum reaction time (3s), indicating that they were behaviourally unresponsive. We took the difference between the moderate and awake reaction time as the measure of the extent that the individuals were affected. Inter-subject similarities were measured as the similarity (Pearson's correlation) of their TTM during the moderate condition. We then correlated the delta reaction time with the inter-subject similarity using Spearman's correlation.

**Table S8: Correlation Inter-subject Similarity of TTM during moderate anaesthesia with delta reaction time (Moderate-Awake). Spearman correlation (Rho) and P-Values.**

| Network Names | Rho | P-Value |
|---|---|---|
| DefaultA 600 | 0.51 | 0.044 |
| DefaultA 800 | 0.33 | 0.213 |
| DefaultB 600 | 0.41 | 0.112 |
| DefaultB 800 | 0.56 | 0.028 |
| DefaultC 600 | 0.1 | 0.705 |
| DefaultC 800 | 0.11 | 0.672 |
| TempPar 600 | -0.09 | 0.746 |
| TempPar 800 | 0.14 | 0.617 |
| ContA 600 | -0.2 | 0.463 |
| ContA 800 | -0.43 | 0.101 |
| ContB 600 | 0.04 | 0.891 |
| ContB 800 | -0.16 | 0.541 |
| ContC 600 | -0.03 | 0.917 |
| ContC 800 | 0.01 | 0.978 |
| LimbicA 600 | 0.08 | 0.771 |
| LimbicA 800 | 0.24 | 0.361 |
| LimbicB 600 | -0.02 | 0.943 |
| LimbicB 800 | 0.13 | 0.633 |
| SalVentAttnA 600 | 0.47 | 0.07 |
| SalVentAttnA 800 | 0.16 | 0.549 |
| SalVentAttnB 600 | 0.35 | 0.188 |
| SalVentAttnB 800 | 0.5 | 0.051 |
| DorsAttnA 600 | -0.59 | 0.019 |
| DorsAttnA 800 | -0.66 | 0.006 |
| DorsAttnB 600 | 0.13 | 0.633 |
| DorsAttnB 800 | 0.01 | 0.969 |
| SomMotA 600 | -0.13 | 0.625 |
| SomMotA 800 | -0.23 | 0.398 |
| Auditory 600 | 0.14 | 0.594 |
| Auditory 800 | 0.02 | 0.952 |
| VisPeri 600 | -0.19 | 0.484 |
| VisPeri 800 | -0.25 | 0.355 |
| VisCent 600 | 0.52 | 0.042 |
| VisCent 800 | 0.36 | 0.165 |
| SUB 54 | -0.03 | 0.926 |

### Supplementary Material 7

**Contrast of awake and deep TTM Shannon entropy during story listening for each network.**

Finally, we investigated whether there were any differences between the awake and deep neural TTMs in terms of the complexity of their distribution. For this we used the built-in MATLAB function "entropy" (bins=256). Below are the results of the contrasts using Wilcoxon signed rank tests.

**Table S7. Wilcoxon signed rank z and p-values of difference in Shannon entropy of TTMs between awake and deep anaesthesia conditions.**

| Network Names | Z-value | P-value |
|---|---|---|
| DefaultA 600 | 2.90 | 0.004 |
| DefaultA 800 | 3.00 | 0.003 |
| DefaultB 600 | 3.00 | 0.003 |
| DefaultB 800 | 3.15 | 0.002 |
| DefaultC 600 | 0.21 | 0.836 |
| DefaultC 800 | 0.52 | 0.605 |
| TempPar 600 | 3.21 | 0.001 |
| TempPar 800 | 3.31 | 0.001 |
| ContA 600 | 1.19 | 0.234 |
| ContA 800 | 1.71 | 0.088 |
| ContB 600 | 0.57 | 0.569 |
| ContB 800 | 1.45 | 0.148 |
| ContC 600 | 2.48 | 0.013 |
| ContC 800 | 2.33 | 0.020 |
| LimbicA 600 | -1.81 | 0.070 |
| LimbicA 800 | -2.17 | 0.030 |
| LimbicB 600 | 0.00 | 1.000 |
| LimbicB 800 | -0.10 | 0.918 |
| SalVentAttnA 600 | 2.12 | 0.034 |
| SalVentAttnA 800 | 2.07 | 0.039 |
| SalVentAttnB 600 | 2.38 | 0.017 |
| SalVentAttnB 800 | 2.17 | 0.030 |
| DorsAttnA 600 | -0.41 | 0.679 |
| DorsAttnA 800 | 0.10 | 0.918 |
| DorsAttnB 600 | 0.26 | 0.796 |
| DorsAttnB 800 | 0.47 | 0.642 |
| SomMotA 600 | -1.76 | 0.079 |
| SomMotA 800 | -1.24 | 0.215 |
| Auditory 600 | -0.26 | 0.796 |
| Auditory 800 | -1.65 | 0.098 |
| VisPeri 600 | -1.40 | 0.163 |
| VisPeri 800 | -2.38 | 0.017 |
| VisCent 600 | -1.50 | 0.134 |
| VisCent 800 | -0.93 | 0.352 |
| SUB 54 | -3.05 | 0.002 |

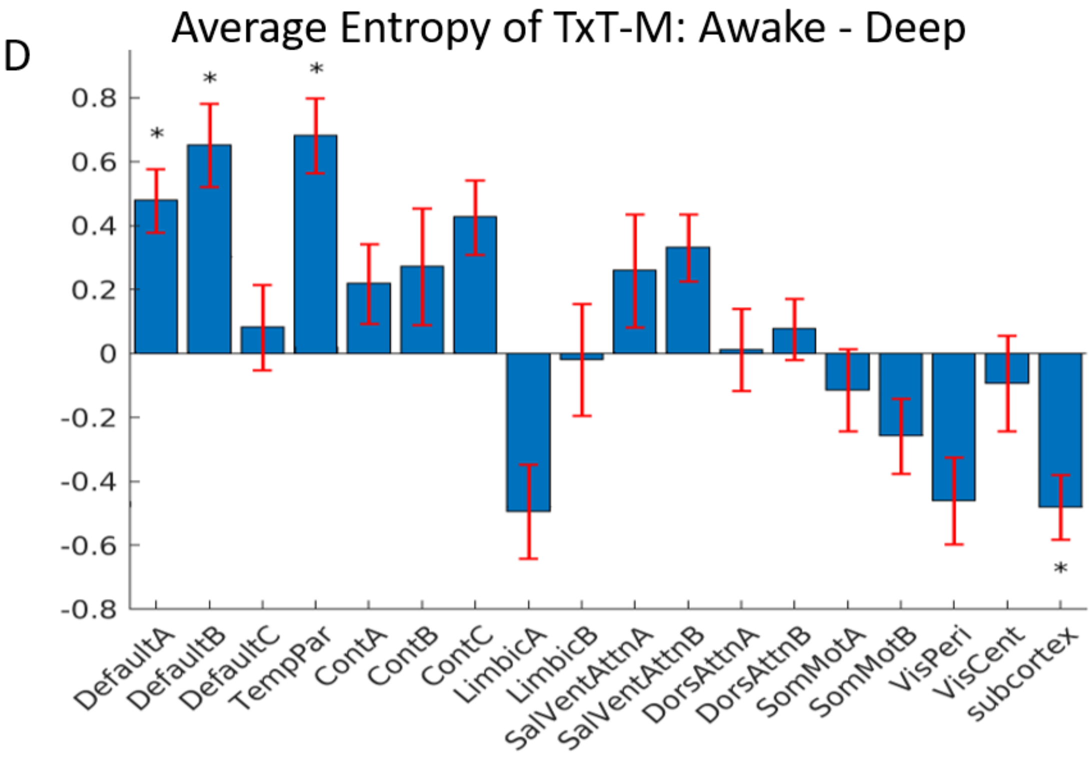


**Figure S6. Shannon entropy differences between awake and deep anaesthesia.**

Average Shannon entropy difference (Awake-Deep) of TTMs for each network. Error bars signify bootstrapped (n=1000) standard error of mean difference. Maps showing which systems significantly display more dissimilarity during consciousness (Deep>Awake; B) and more inter-subject similarity (IntSS) during consciousness (Awake>Deep; C). Asterisks indicate FEW corrected significances. Error bars indicate bootstrapped (n=1000) standard error of mean difference.

Note, for the correlation between the difference in intersubject similarity between consciousness and unconsciousness and the difference between complexity of the TTMs in consciousness and unconsciousness, we used all available networks (i.e., 18), as presented throughout the supplementary materials. The reason for this is that this would give a more reliable statistical estimate, instead of using the 4 networks that are reported in the main text.

### Supplementary Material 8

**Summary of results and supplementary discussion to the results not discussed in the main text.**

We here review all results together to assess other possible neural correlates of general and special experiences during story listening. The are many results, and with several methodological contingencies (see methods) which we explored extensively (e.g., Supplementary materials 1, 2, and 3). Therefore, we have placed a summary of all the results in table S10. This shows analyses (columns) results for each network (Rows), and indicates with a tick (✓) If there was strong evidence, and a tilde (~) for moderate evidence for that analysis across analytical contingencies (see methods and supplementary materials for details).

In the supplementary discussion below, we will review each network that is proposed in the neural correlates of general and special experience (table S9) in relation to previous research.

**Table S9. Summary of results for each network across all analyses (see methods and supplementary materials) and inclusion into proposed neural correlates of general or special experience.** The columns represent different analyses, presented above; whilst the rows networks (n=18). The final column represents whether the network was included into the neural correlates of general or special experience. A tick (✓) signifies strong convergence of evidence across analyses (See methods and supplementary materials) whilst a tilde (~) represents moderate evidence. Empty cells represent little or no evidence in specific analysis (corresponding to column). Networks and parcellations have been previously described (A. Schaefer et al., 2018; Yeo et al., 2011). In parentheses are the relevant supplementary materials for each type of analysis.

| | **Subjective ∝ Neural Correspondence (average) (S2)** | **Subjective ∝ Neural correspondence Variation (S3)** | **RT ∝ Inter-Subject Similarity Breakdown (S8)** | **Story specific effects (Interaction with rest) (S4)** | **Greater or lower IntSS in consciousness. (S6)** | **Possible neural correlates of general and special experience** |
|---|---|---|---|---|---|---|
| **DMN A** | ✓ | ✓ | ~ | | ✓ | **SPECIAL** |
| **DMN B** | ✓ | | ~ | ✓ | ✓ | **SPECIAL** |
| **DMN C** | | ~ | | | ✓ | |
| **TempPar** | | | | ✓ | ✓ | **SPECIAL** |
| **FPN A** | ✓ | ~ | | | | |
| **FPN B** | | ✓ | | | | |
| **FPN C** | ✓ | | | ✓ | ✓ | **SPECIAL** |
| **LIN A** | | | | ✓ | ✓ | **GENERAL** |
| **LIN B** | | | | | ✓ | |
| **VAN A** | ✓ | ✓ | | | ✓ | **SPECIAL** |
| **VAN B** | | | | ✓ | ✓ | **SPECIAL** |

| | | | | | | |
|---|---|---|---|---|---|---|
| **DAN A** | ~ | ~ | ✓ | | ~ | **GENERAL** |
| **DAN B** | ✓ | ✓ | | | | |
| **SM A** | ✓ | ✓ | | ✓ | | **GENERAL** |
| **AUD** | ✓ | ✓ | | ✓ | ✓ | **GENERAL** |
| **VPN** | ✓ | | | | ✓ | **GENERAL** |
| **VCN** | | ~ | | ✓ | ✓ | **SPECIAL** |
| **SUB** | ~ | ~ | | | ✓ | **GENERAL** |

**Supplementary discussion.**

In this supplementary section, we review the results of the networks not discussed in the main text in relation to previous results.

Our main objective was to propose preliminary neural correlates of general and special experience. Given our exploratory results, evidencing involvement in the task and higher inter-subject dissimilarity in consciousness, we suggest that, beyond the DMN, the ventral attention network (VAN) dynamics are involved in individual specific experience (For reference see figures S2, S3, S4 and S5; or table S10). Furthermore, it seems that a precuneal component of the frontalparietal control network (FPN-C) and the early visual network also display individual-specific dynamics. Conversely, given relevance to the task and higher similarity in consciousness, the dynamics of the subcortex, the visual peripheral network, the somato-motor network and Limbic A all seem to support relatively more generalisable streams of consciousness across the different individual experiences. We shall now briefly discuss these networks in relation to previous literature.

Adjacent to the posterior DMN-A regions, the FPN-C also includes a dorsal precuneal and a ventral posterior cingulate region (sometimes included in the DMN (Cavanna, 2007; Lyu et al., 2021). These precuneal and cingulate regions are thought to compose an integratory hub supporting the most complex cognition (Cavanna, 2007; Leech et al., 2012; Lyu et al., 2021; Margulies et al., 2016; van den Heuvel & Sporns, 2011). Such posterior regions have also been found to track the narrative of stories at their most abstract levels, so much so that they are generalisable across modalities (Baldassano et al., 2017; Chen et al., 2017). This network was found to track the average of the subjective TTM and to display a differential effect between rest and story listening as a function of (un)consciousness (S2, S4 and Table S10).

The temporal parietal (TempPar) network was originally included in the DMN (in 7 network definition as opposed to the 17 network definition) and comprises superior temporal and parietal regions (angular gyrus specifically). These, similarly to the DMN-B discussed in the main text, are regions commonly associated with language related processing. In fact, when compared to neurosynth meta-analytic maps, this network had a substantial overlap with maps associated with the term “language” (57.6 %) and “semantic” (36%). Thus, broadly speaking, interpretations are equivalent to those proposed for the DMN-B in the main text. However, see text at the end of supplementary materials 2 and 3 for additional interesting findings regarding the TemPar network (i.e., different behaviour in rest and auditory story conditions, negative correlations with behavioural rating once autocorrelation is removed).

Conversely to the DMN -B and the TempPar, the limbic A network is also comprised of temporal regions (inferior and anterior pole), but is seemingly involved in common aspects of experience. Also a higher order network (Margulies et al., 2016) these regions are known to be involved in object recognition, semantic processing and social cognition (Conway, 2018; Lambon Ralph et al., 2016; Pehrs et al., 2015), which are known to be affected by propofol sedation (Adapa et al., 2014). Thus, whilst the inferior temporal ventral stream, known to be involved in object recognition (e.g., (Conway, 2018; Pehrs et al., 2015) may underlie common experiences, the more middle temporal and angular gyri (DMN-B and TempPar) may underlie more individual-specific semantically and mnemonically driven experiences (Chen et al., 2017; Friederici, 2011; Hagoort, 2019; Smallwood et al., 2016).

Of less easy interpretation is the involvement in the ventral attention network (VAN; also called salience in (A. Schaefer et al., 2018; Yeo et al., 2011). Interpreted as being involved in stimulus-driven attention (Vossel et al., 2014), the VAN-A’s dynamics has shown correlations to suspense dynamics (fig. S2), but also the tracking of individual differences, second only to the DMN-A (fig. S3). The VAN B instead showed to behave differently in the story versus rest conditions, showing virtually no effect in IntSS resting-state effects across levels of anaesthesia, but a substantial scaling during story listening (fig. S4). Although “stimulus-driven attention” would indicate common experience (the stimulus being the commonality), what counts as a “salient” stimulus shows very large and consistent individual differences (De Haas et al., 2019). Furthermore, this network’s dynamics may reportedly also be driven by “internal” memory-driven stimuli, (Corbetta et al., 2008; Seeley, 2019; Vossel et al., 2014), which will be specific to the individual’s development. In fact pathological (Seeley, 2019) and personality differences ((Tian, Wang, Xu, Li, & Ma, 2018) have been found in relation to this network, indicating that it may underlie a substantial portion of individual differences.

Thus, beyond the classic DAN-DMN antithesis discussed in the main text (Huang et al., 2020), another dynamic balance may underlie individual and common experiences: The ventral and dorsal attention networks respectively. These networks, described as being circuit breakers for one another (Corbetta et al., 2008) engage in a dynamic and flexible interplay (Vossel et al., 2014), that would tendentially support special and common attention-driven experiences.

Conversely to the auditory network reported in the main text, the correlation with suspense dynamics of story with visual and somato-motor dynamics perhaps may be counterintuitive (fig. 1I, fig. 2D). However, it has been repeatedly shown that information in one sensory modality may instigate processing in other sensory modalities (Adolphs et al., 2000; Akama et al., 2012; De Borst & De Gelder, 2017; Pulvermuller, 2005; Sanchez et al., 2020; M. Schaefer et al., 2015; Vetter et al., 2014). The complete irrelevance of the visual and motor system in representing auditory information is unlikely. For example, visual peripheral network (VPN) dynamics are known to be primed by attention/stimuli-salience in a behaviourally relevant way (Cate et al., 2009) as well as it's dynamics potentially being relevant to content imagery (Vetter et al., 2014), which may be of particular interest to the auditory story condition.

The involvement of the SMN-A may also be controversial. However its emotional processing in naturalistic social settings (Adolphs et al., 2000; Kropf et al., 2019; M. Schaefer et al., 2015) and in auditory imagery (De Borst & De Gelder, 2017) is well documented, suggesting a role in embodied representations. In fact, several bodily actions are inferable from the auditory story presented to participants (S10). The Visual Central Network, is a surprising finding for the neural correlates of special experience (see table S9). One potential explanation is differences between individual's use of the early visual cortex during an auditory story (Reeder, 2017; Vetter et al., 2014). Differences may lie in imagery vividness, content of visualisation and visual/behavioural priming strategies (e.g., (Cate et al., 2009; Reeder, 2017; Zeman, Dewar, & Della Sala, 2015). Nonetheless, its involvement in the task is evidenced by the interaction analysis (fig. S4) and its correlation with the average and the variance of the subjective dynamics (fig. S2 and S3).

Finally, The subcortex (excluding cerebellum, described in methods section)is the oldest and physiologically most fundamental network to be included in this study (Panksepp, 2011). It is well suited to underlie common experiences in that it supports survival related processes. In fact, we found that it's dynamics may track that of subjective feelings of suspense and also that its variations across individuals may potentially track individual differences in subjective reporting (Although note, only with the deconvolution method rather than the lagged method, see S2). Although our data (fig. S5) suggests that the subcortical dynamics are more similar in consciousness compared to unconsciousness, we cannot discount experience-driven individual differences in subcortical processing (e.g., Péron, Frühholz, Ceravolo, & Grandjean, 2015; Sylvester et al., 2020; Telzer et al., 2013). An interesting finding, is that the subcortex is the only network that consistently displayed increased complexity in the unconscious condition (fig. S6), perhaps indicating a disorganisation in dynamics.

Finally, although our analyses permitted a relative categorisation of a network as underlying general or special experiences, this coarse distinction is likely to be revealed as unimaginably more complex as new techniques and paradigms permitting a concurrent analysis at multiple dimensions (experiential, spatial and temporal) are developed (note, this is copied from main text). This may partly explain why the FPN-A, despite being involved in the task and showing a trend in heightened similarity in consciousness, did not display sufficient evidence to be included in general neural correlates of experience (Naci et al., 2017, 2018).

Another interesting finding in relation to this is that the network dynamics that display increased individual differences in consciousness tend to contain more information (as measured by Shannon entropy), whilst the dynamics that are more similar between individuals in consciousness are relatively more structured. Increased information of intrinsic dynamics would naturally result in less inter-subject similarity. However, this finding may also elucidate the dynamic equilibrium between more personal-subjective and more communicable-objective processes that the brain has to constantly negotiate (Carhart-Harris & Friston, 2010; Friston, 2010). Presumably, certain pathologies may be partially explained by a deviation of such an equilibrium (Carhart-Harris & Friston, 2019; Froggatt, 2005; Fromm, 1998). Further evidencing the involvement of DMN regions in special experience is that these networks are the only ones showing a more "informative" distribution of self-similarities in consciousness during story listening (fig. 3D), mirroring theoretical predictions

(Carhart-Harris et al., 2014). In fact, in line with typical notions of consciousness (Carhart-Harris et al., 2014) the control awake condition tended to show more entropy over all networks (fig. S6). Only four networks showed a family-wise error corrected effect across granularity. The DMN-A, DMN-B and TemParnetworks showed higher entropy during the consciousness condition, whilst the subcortex showed the opposite effect (see S7 for full results).

Another apparently contradictory result in relation to previous work, is the higher subcortical dynamics complexity (measured via Shannon Entropy) in unconsciousness. Previous results (Coppola et al., 2022) show heightened complexity of subcortical dynamics by measuring the temporal information of various graph theory properties and intrinsic dynamics in rest. However, while these papers measured complexity of temporal sequences, here we applied Shannon entropy looking at the complexity of the distribution of intrinsic similarity (which is shown to diminish in consciousness in proximal transitions, Coppola et al., in press). Thus, here we discard temporally-specific information of moment-to-moment changes and focus on the absolute variation of intrinsic similarity values. This suggests that despite having a higher complexity in temporal sequences in consciousness (resting state), the subcortex may display a more organised structure in the long-term temporal distribution during such a story (S9; e.g., visiting similar "states", or travelling similar distances overtime), which may translate to a particularly predictive desynchronisation in unconsciousness  (Tsurugizawa & Yoshimaru, 2021).
